# FGF21 Counteracts Alcohol Intoxication by Activating Noradrenergic Neurons

**DOI:** 10.1101/2022.08.09.502667

**Authors:** Mihwa Choi, Marc Schneeberger, Wei Fan, Abhijit Bugde, Laurent Gautron, Kevin Vale, Yuan Zhang, Jeffrey M. Friedman, David J. Mangelsdorf, Steven A. Kliewer

## Abstract

Animals that consume fermenting fruit or nectar are exposed to ethanol, thus increasing their risk of injury or predation. This risk is heightened in humans, who have actively imbibed alcohol for thousands of years. In this report, we show that the hormone FGF21, which is strongly induced by ethanol in murine and human liver, exerts sobering or “amethystic” effects on both arousal and motor coordination without changing ethanol catabolism. Mice lacking FGF21 take longer than wild-type littermates to recover their righting reflex and balance following ethanol exposure. Conversely, pharmacologic FGF21 administration reduces the time needed for mice to recover from ethanol-induced unconsciousness and ataxia. FGF21 mediates it amethystic effects by directly activating the noradrenergic nervous system, which regulates arousal and alertness. These results indicate that this FGF21 liver-brain pathway evolved to protect against ethanol-induced intoxication and that it might be targeted pharmaceutically for treating acute alcohol poisoning.

## INTRODUCTION

Simple sugars in ripening fruits and nectars are a rich source of calories for many animals. However, consumption of ethanol produced by the natural fermentation of these sugars can cause intoxication, thus impairing mobility and judgement (Dudley and Maro, 2021).Accordingly, animals have evolved enzymes that efficiently catabolize ethanol. Humans, who actively ferment and distill ethanol, are particularly susceptible to the detrimental effects of acute alcohol overconsumption, which include dehydration, disorientation and loss of consciousness, impaired breathing and heart function, and even death. Emergency treatment for alcohol poisoning includes keeping intoxicated individuals upright and awake, if possible. Thus, drugs with sobering or “amethystic” (anti-intoxicant) activity could prove valuable for treating this dangerous condition (Alkana and Noble, 1979)

FGF21 is a hormone that is induced in liver by a variety of metabolic stresses including starvation, protein deficiency, simple sugars and ethanol (Fisher and Maratos-Flier, 2016; Owen et al., 2015). In humans, ethanol is by far the most potent inducer of FGF21 described to date (Desai et al., 2017; Søberg et al., 2018; Song et al., 2018). Previous studies showed that FGF21 suppresses ethanol preference (Flippo et al., 2022; Schumann et al., 2016; Talukdar et al., 2016), induces water drinking (Song et al., 2018), and protects against alcohol-induced liver injury (Desai et al., 2017; Liu et al., 2016; Zhu et al., 2014). Thus, FGF21 plays a broad role in defending against the harmful consequences of ethanol exposure.

FGF21 acts on a heteromeric cell surface receptor composed of a conventional FGF receptor tyrosine kinase (FGFR1c) in complex with the single-pass transmembrane protein, β-Klotho (KLB) (Ogawa et al., 2007). FGF21 binds directly to both FGFR1c and KLB, with FGFR1c serving as the downstream signaling effector. Human genome-wide association studies have linked SNPs in and around both the *FGF21* and *KLB* genes to increased alcohol consumption (Frayling et al., 2018; Schumann et al., 2016; Søberg et al., 2017), further underscoring the important relationship between FGF21 and ethanol. In mice, FGF21 acts on its receptor complex in the nervous system both to suppress ethanol preference and to induce water consumption (Schumann et al., 2016; Song et al., 2018).

Norepinephrine (NE) is an abundant neuromodulator in the CNS. Most central NE is synthesized in the locus coeruleus (LC), a small nucleus in the pons of the brainstem. LC neurons project extensively throughout the brain to regulate diverse biological processes, including arousal and alertness (Berridge et al., 2012; Poe et al., 2020). In this report, we show that FGF21 directly activates noradrenergic neurons in the LC region. We further examine the consequences of inducing this FGF21-NE pathway in the context of ethanol-induced loss of balance and impaired motor coordination. These results show that FGF21 is an endogenous amethystic agent.

## RESULTS

### FGF21 deficiency exacerbates ethanol-induced intoxication

We administered a single, binge dose of ethanol (5 g/kg by oral gavage) to wild-type (WT) and global *Fgf21*^*–/–*^ mice. Consistent with prior results, plasma FGF21 was induced by ethanol in WT mice, peaking at 2 hours (Figure 1A). We next examined the animals’ righting reflex, a standard marker of inebriation. While both WT and *Fgf21*^*–/–*^ mice lost their righting reflex ∼30 minutes after ethanol gavage (Figure 1B), *Fgf21*^*–/–*^ mice required several more hours than WT mice to recover (Figure 1C). WT and *Fgf21*^*–/–*^ mice cleared ethanol from the plasma at the same rate (Figure 1D), and brain ethanol concentrations were similar between the genotypes (Figure 1E). Thus, FGF21 protects against ethanol-induced loss of righting reflex without affecting ethanol catabolism.

**Figure 1.**
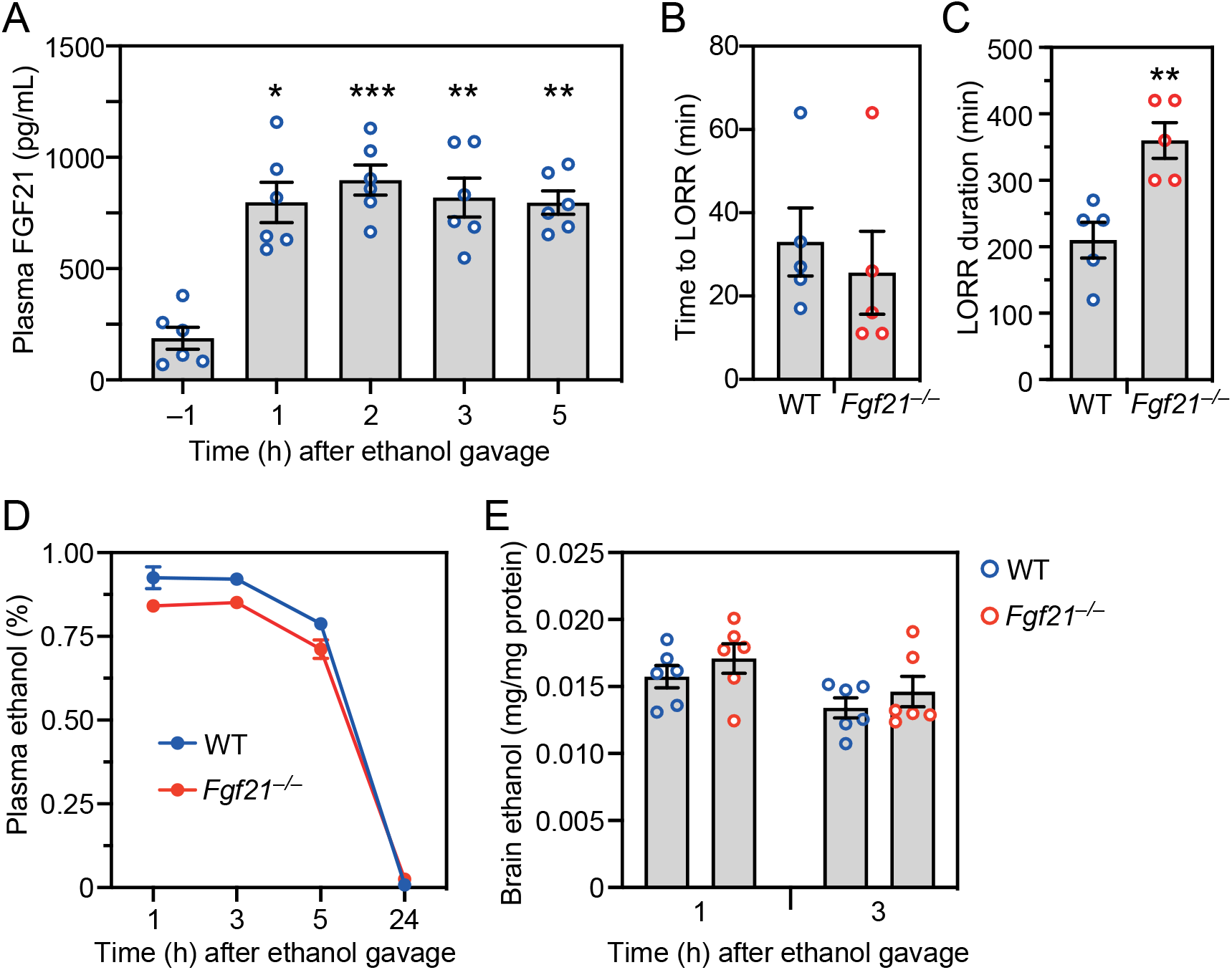
*Fgf21*^−/−^ mice have a prolonged righting reflex recovery time after a binge ethanol dose. (A) Plasma FGF21 concentrations were measured in wild-type (WT) mice either 1 hour before administering ethanol (5 g/kg, oral gavage) or at the indicated times after ethanol (n = 6 mice/group). (B-E) WT and *Fgf21*^**−/−**^ mice were administered ethanol (5 g/kg, oral gavage) and the following measurements made: time to loss of righting reflex (LORR) (B); LORR duration (C); plasma ethanol concentrations (D); brain ethanol concentrations (E) (n = 5-6 mice/group). All data represent the mean ± SEM. *, *P* < 0.05, **, *P* < 0.01, ***, *P* < 0.001 compared to -1 hour (A) or WT (C).

We obtained similar results for both time to loss of righting reflex and its duration with hepatocyte-specific *Fgf21*-knockout (*Fgf21*^*Alb*^) mice (Figures 2A and 2B) and neuron-specific *Klb*-knockout (*Klb*^*Camk2a*^) mice (Figures 2D and 2E). There were no differences in plasma ethanol clearance between the knockout lines and control mice (Figures 2C and 2F). These results indicate that liver-derived FGF21 attenuates ethanol-induced loss of righting reflex by acting on its receptor in the nervous system.

**Figure 2.**
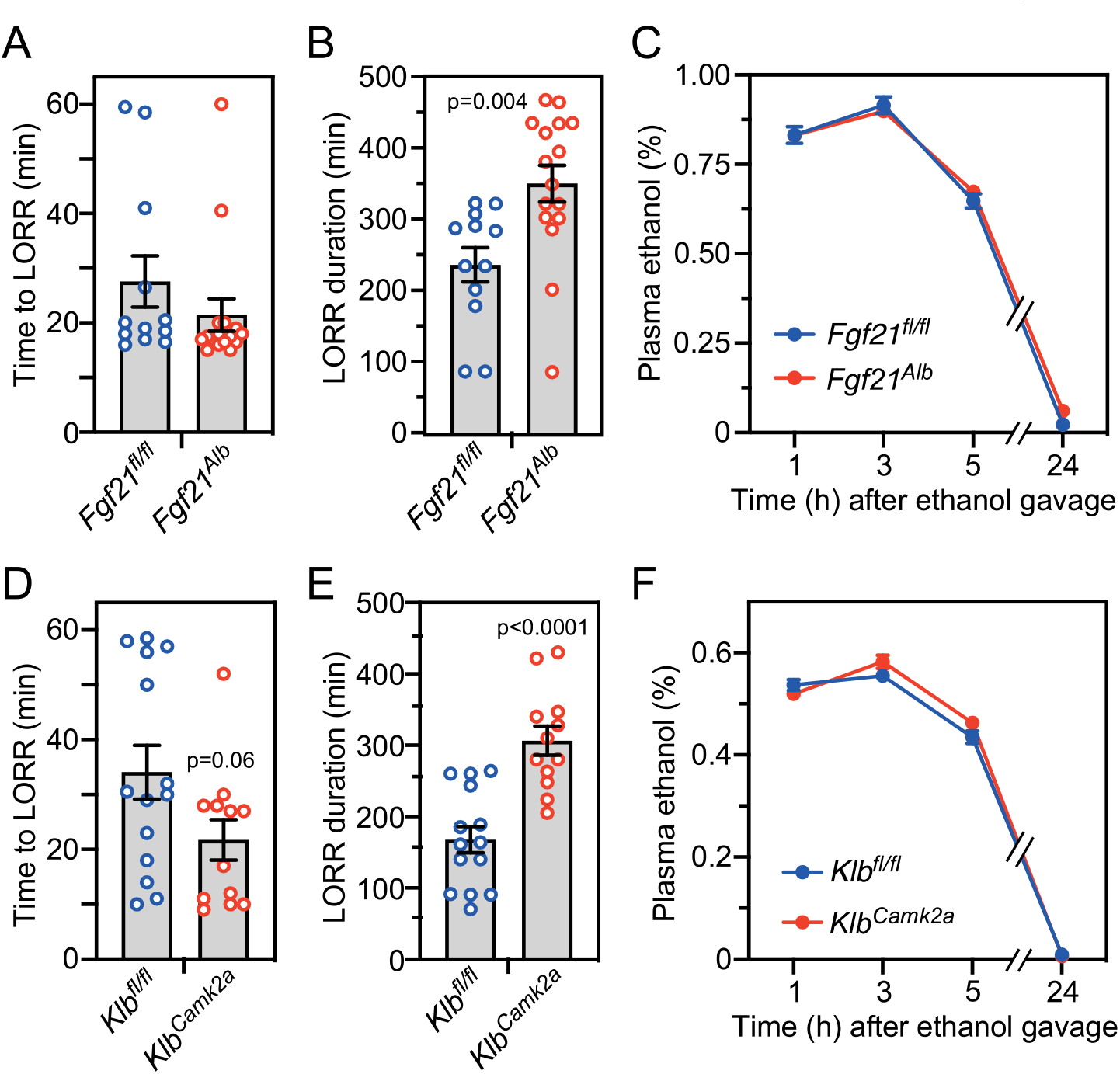
Hepatocyte-specific *Fgf21*^−/−^ and neuron-specific *Klb*^−/−^ mice have prolonged righting reflex recovery times after a binge ethanol dose. Control (*Fgf21*^*fl/fl*^) and hepatocyte-specific *Fgf21*^−/−^ (*Fgf21*^*Alb*^) mice (A-C) or control (*Klb*^*fl/fl*^) and neuron-specific *Klb*^−/−^ (*Klb*^*Camk2a*^) mice (D-F) were administered ethanol (5 g/kg, oral gavage) and the following measurements made: time to loss of righting reflex (LORR) (A, D); LORR duration (B, E); and plasma ethanol concentrations (C, F) (n = 12-16 mice/group for A, B, D, E and 6 mice/group for C and F). All data represent the mean ± SEM.

### Pharmacologic administration of FGF21 is amethystic

We next examined whether pharmacologic FGF21 treatment of WT mice decreases the time to righting reflex recovery after ethanol administration. WT mice were administered a high ethanol dose (5 g/kg) by oral gavage followed by i.p. FGF21 injection one hour later.

Remarkably, FGF21 administration reduced the time required for both male and female mice to recover their righting reflex by ∼90 minutes, reflecting a roughly 50% decrease compared to vehicle-treated mice (Figures 3A and 3B). This effect was dose dependent and maximally efficacious at 1 mg/kg FGF21 in male mice (Figure 3A).

**Figure 3.**
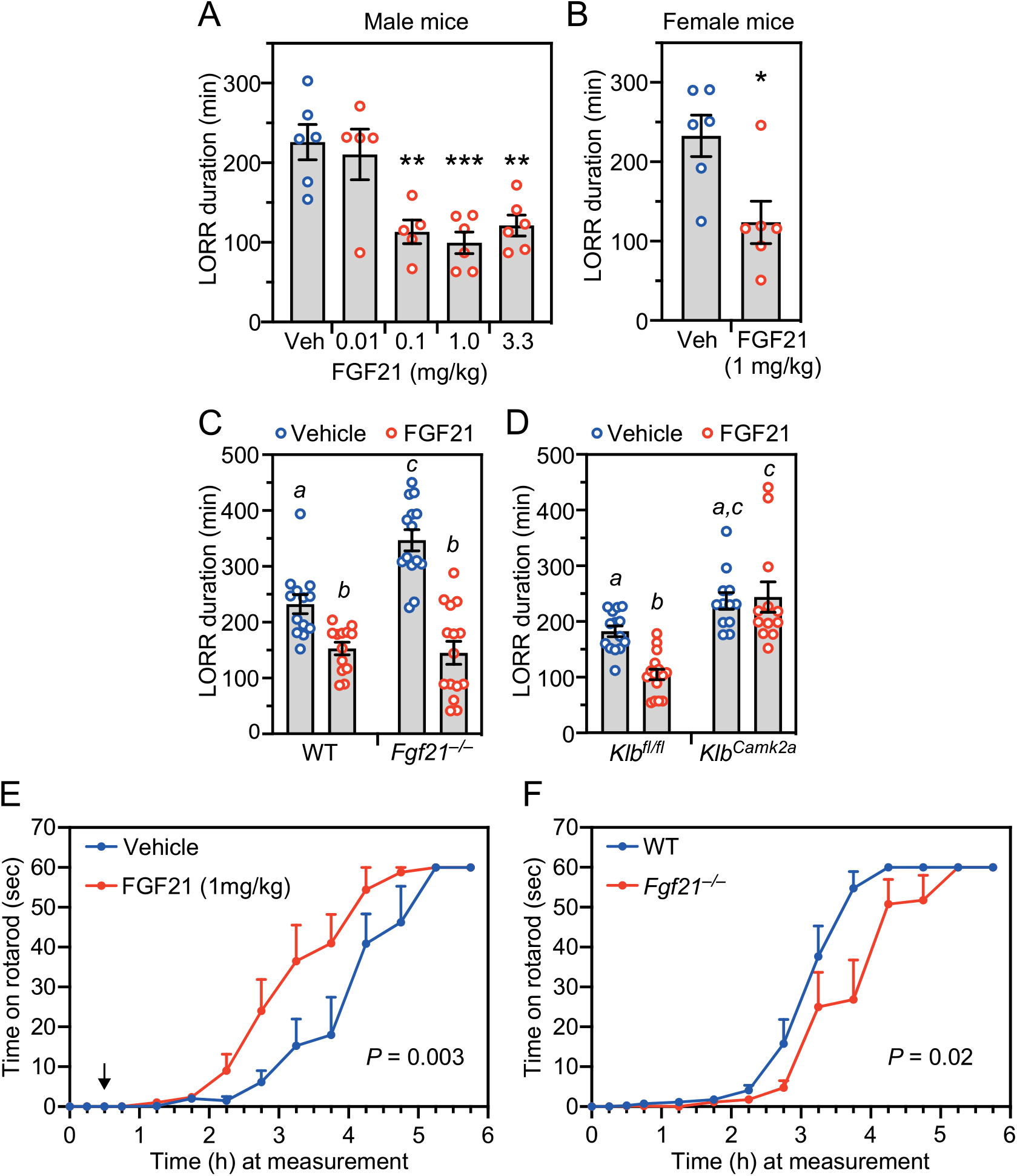
Pharmacologic FGF21 attenuates alcohol-induced loss of righting reflex and ataxia. (A, B) Wild-type (WT) male (A) or female (B) mice were administered ethanol (5 g/kg, oral gavage) followed 1 hour later by i.p. injection of vehicle or FGF21 at the indicated doses. Loss of righting reflex (LORR) duration was measured after FGF21 or vehicle administration (n = 5-6 mice/group). Data represent the mean ± SEM. *, *P* < 0.05, **, *P* < 0.01, ***, P < 0.001 compared to vehicle. (C, D) WT and *Fgf21*^**−/−**^ mice (n = 13-15 mice/group) (C) or control (*Klb*^*fl/fl*^) and neuron-specific *Klb*^**−/−**^ (*Klb*^*Camk2a*^) mice (n = 12-16 mice/group) (D) were administered ethanol (5 g/kg, oral gavage) followed 1 hour later by i.p. injection of FGF21 (1 mg/kg) or vehicle. LORR duration was measured after FGF21 or vehicle administration. Data represent the mean ± SEM. Different lowercase letters indicate statistical significance (*P* < 0.05). (E)WT mice were administered ethanol (2 g/kg, i.p.) followed 30 minutes later by injection of FGF21 (1 mg/kg, i.p.; indicated by arrow) or vehicle. The time mice could remain on a spinning rotarod was measured, with 60 seconds the maximum (n = 8 mice/group). (F)WT and *Fgf21^−/−^* mice were administered ethanol (2 g/kg, i.p.) and the time they could remain on a spinning rotarod measured as in (E) (n = 6-7 mice/group). All data represent the mean ± SEM.

We performed this same pharmacologic FGF21 rescue experiment in *Fgf21*^*−/−*^ and *Klb*^*Camk2a*^ mice. In *Fgf21*^*−/−*^ mice, FGF21 administration reduced the righting reflex recovery time to that seen in WT mice (Figure 3C). In contrast, FGF21 had no effect on righting reflex recovery time in *Klb*^*Camk2a*^ mice (Figure 3D), indicating that pharmacologic FGF21 exerts its amethystic effect via the nervous system.

We also investigated whether FGF21 inhibits ethanol-induced impairment of motor coordination. Pharmacologic FGF21 treatment reduced the time required for WT mice to recover their coordination on a rotarod following administration of a moderate dose of ethanol (2 g/kg, i.p.) (Figure 3E). Conversely, recovery time was significantly increased in *Fgf21*^*−/−*^ compared to WT mice (Figure 3F). Thus, FGF21 is amethystic for ethanol-induced impairment as measured by both righting reflex and rotarod performance.

### FGF21’s amethystic activity is selective for ethanol

We next tested whether FGF21 counteracts other sedatives, including the glutamatergic receptor antagonist ketamine and the GABA receptor agonists diazepam and pentobarbital. Because these sedatives act more quickly and for a shorter duration than ethanol, we compressed the experimental timeline: mice were i.p. injected with either ethanol or each of the other sedatives followed by i.p. injection of FGF21 or vehicle 30 minutes later. FGF21 retained its ability to reduce righting reflex recovery time in ethanol-treated mice under these modified conditions (Figure 4A). In contrast, FGF21 administration did not reduce the righting reflex recovery time for ketamine, diazepam and pentobarbital (Figures 4B-D). Thus, FGF21’s amethystic activity is selective for ethanol relative to these other drugs.

**Figure 4.**
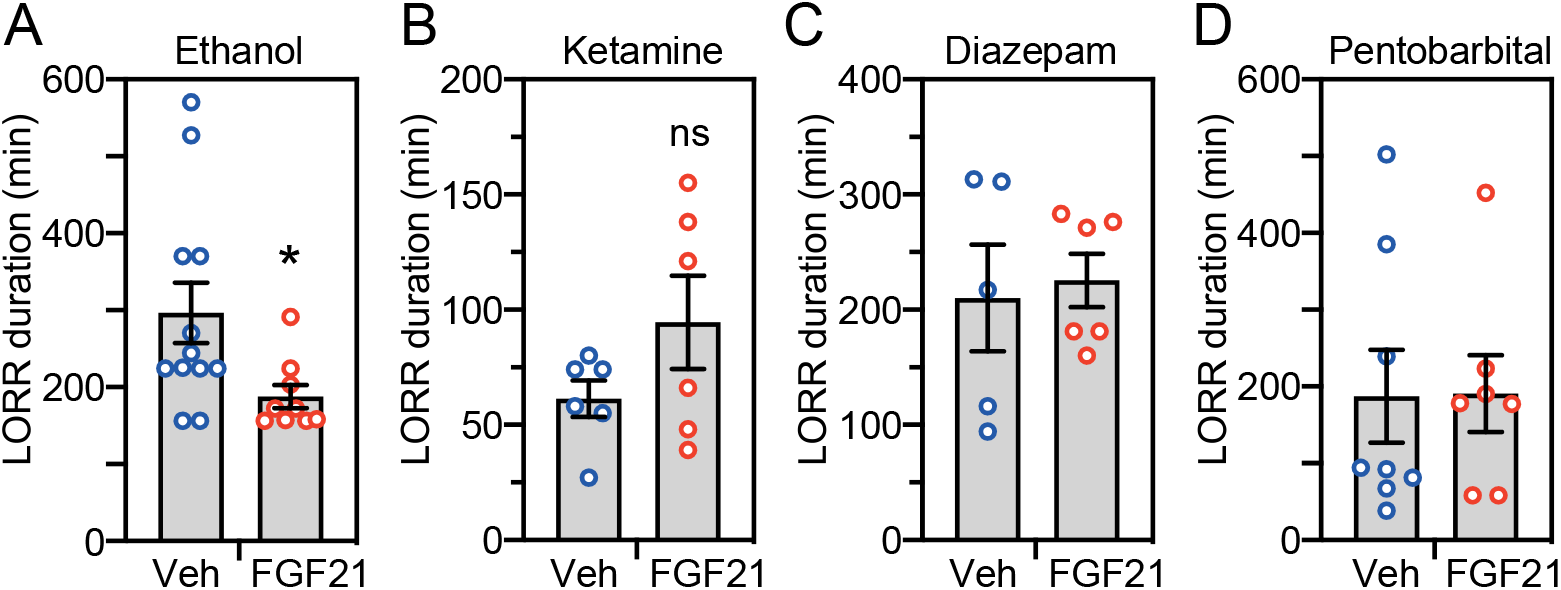
Pharmacologic FGF21 does not attenuate other sedatives. Wild-type mice were administered ethanol (4.3 g/kg) (A) (n = 9-12 mice/group), ketamine (200 mg/kg) (B) (n = 6 mice/group), diazepam (30 mg/kg) (C) (n = 5-6 mice/group) or pentobarbital (55 mg/kg) (D) (n = 7-8 mice/group) by i.p. injection. After 30 minutes, mice were i.p. injected with either FGF21 (1 mg/kg) or vehicle. Loss of righting reflex (LORR) duration was measured after FGF21 or vehicle administration. All data represent the mean ± SEM. *, *P* < 0.05 compared to vehicle-treated mice.

### FGF21 is a physiologic regulator of noradrenergic neurons

In a previous study, it was shown that dopamine β-hydroxylase (*Dbh*)-KO mice, which are unable to synthesize NE, have a prolonged righting reflex recovery time in response to ethanol without any change in ethanol catabolism (Weinshenker et al., 2000), similar to the responses we observed in *Fgf21*^*−/−*^ mice. Moreover, studies in mice and rats showed that ethanol administration activates neurons in the LC, the principal site of NE synthesis (Chang et al., 1995; Kolodziejska-Akiyama et al., 2005; Thiele et al., 1997). These findings led us to investigate whether physiologic FGF21 is responsible for ethanol-induced activation of noradrenergic neurons in the LC. WT and *Fgf21*^*−/−*^ mice were administered ethanol (5 g/kg) by oral gavage and sacrificed 2.5 hours later. Immunostaining of LC sections was performed for c-Fos and the NE transporter (NET), which are markers of neuronal activity and noradrenergic neurons, respectively. As previously reported, ethanol induced c-Fos expression in NET^+^ LC neurons of WT mice (Figures 5A and 5B) (Chang et al., 1995; Kolodziejska-Akiyama et al., 2005; Thiele et al., 1997). Remarkably, this effect was completely absent in *Fgf21*^*−/−*^ mice (Figures 5A and 5B). The total number of NET^+^ cells was equivalent in WT and *Fgf21−/−* mice indicating that the absence of FGF21 does not compromise the neuronal lineage (Figure 5B). These data demonstrate that FGF21 is required for ethanol to activate noradrenergic neurons in the LC.

**Figure 5.**
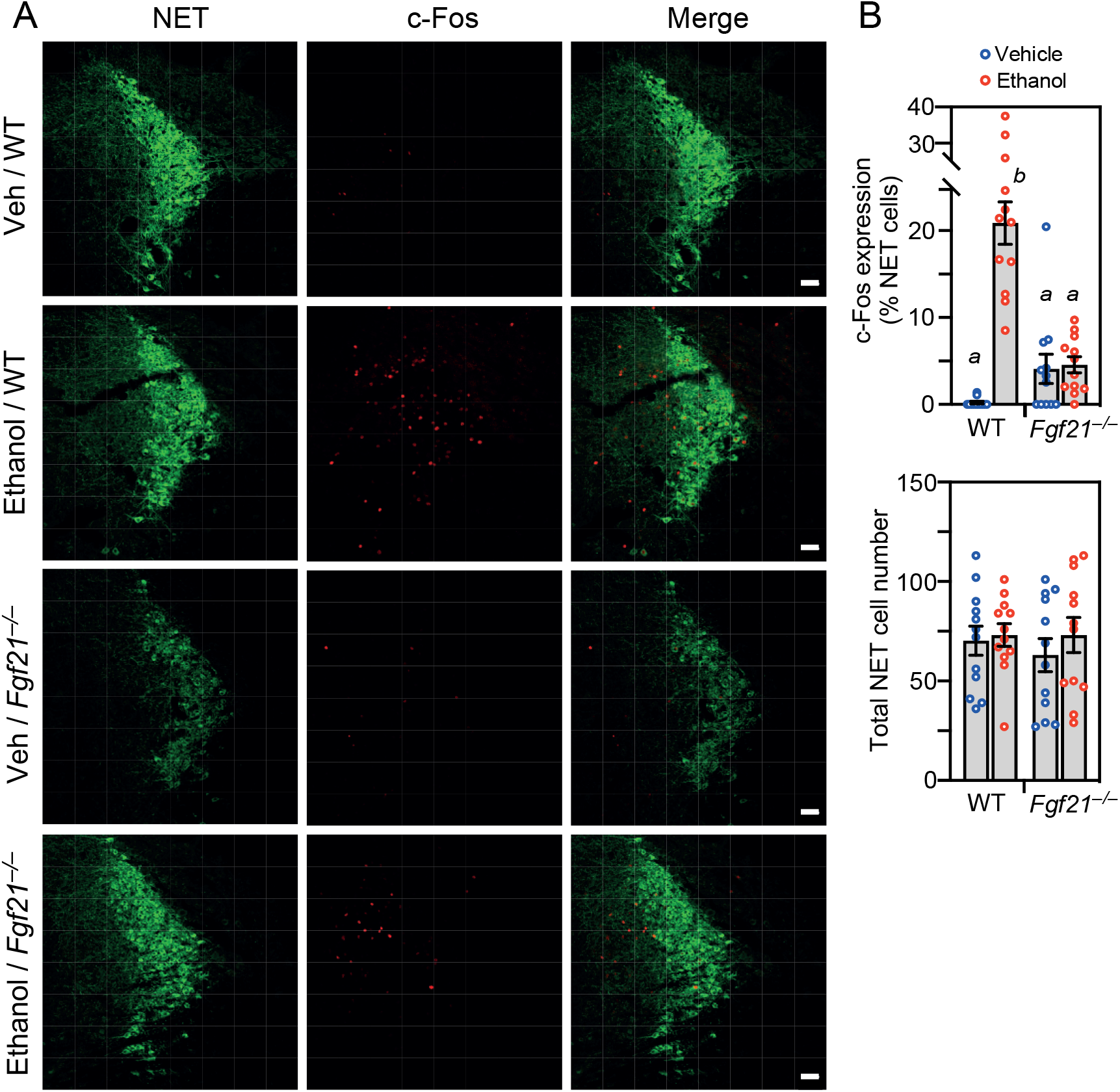
FGF21 is a physiologic regulator of noradrenergic neurons. Representative confocal images of immunostaining for c-Fos and norepinephrine transporter (NET) in locus coeruleus sections prepared from wild-type (WT) and *Fgf21*^**−/−**^ mice 2.5 hours after oral gavage with either water or ethanol (5 mg/kg). Scale bars represent 50 μM. Quantification of c-Fos/NET co-expression (upper panel) and total NET-positive cell number (lower panel) (n = 3 sections/mouse, 4 mice/group). Data represent the mean ± SEM. Different lowercase letters indicate statistical significance (P < 0.05).

### FGF21 acts directly on noradrenergic neurons

To determine whether FGF21 acts directly on noradrenergic neurons, we first examined whether FGF21’s obligate co-receptor, KLB, which is selectively expressed in only a few regions of the brain (Bookout et al., 2013), is present in the LC. Due to the lack of reliable KLB antibodies, we measured KLB expression using an established reporter mouse in which tdTomato was knocked into the endogenous *Klb* gene, resulting in a KLB-tdTomato fusion protein (Coate et al., 2017). The fusion protein was detected by immunostaining in the LC, where it co-localized with NET in most but not all cells (Figure 6A). The fusion protein was also detected in regions adjacent to the LC where there was little or no NET staining (Figure 6A). Thus, KLB is expressed in noradrenergic neurons and other cell types in and around the LC. Due to the lack of antibodies selective for FGFR1c, which is the second constituent of the FGF21 receptor complex, we examined its expression at the mRNA level by in situ hybridization. *Fgfr1c* mRNA was broadly expressed throughout the LC region, where it colocalized with *Klb* mRNA in some noradrenergic neurons (Figure S1).

**Figure 6.**
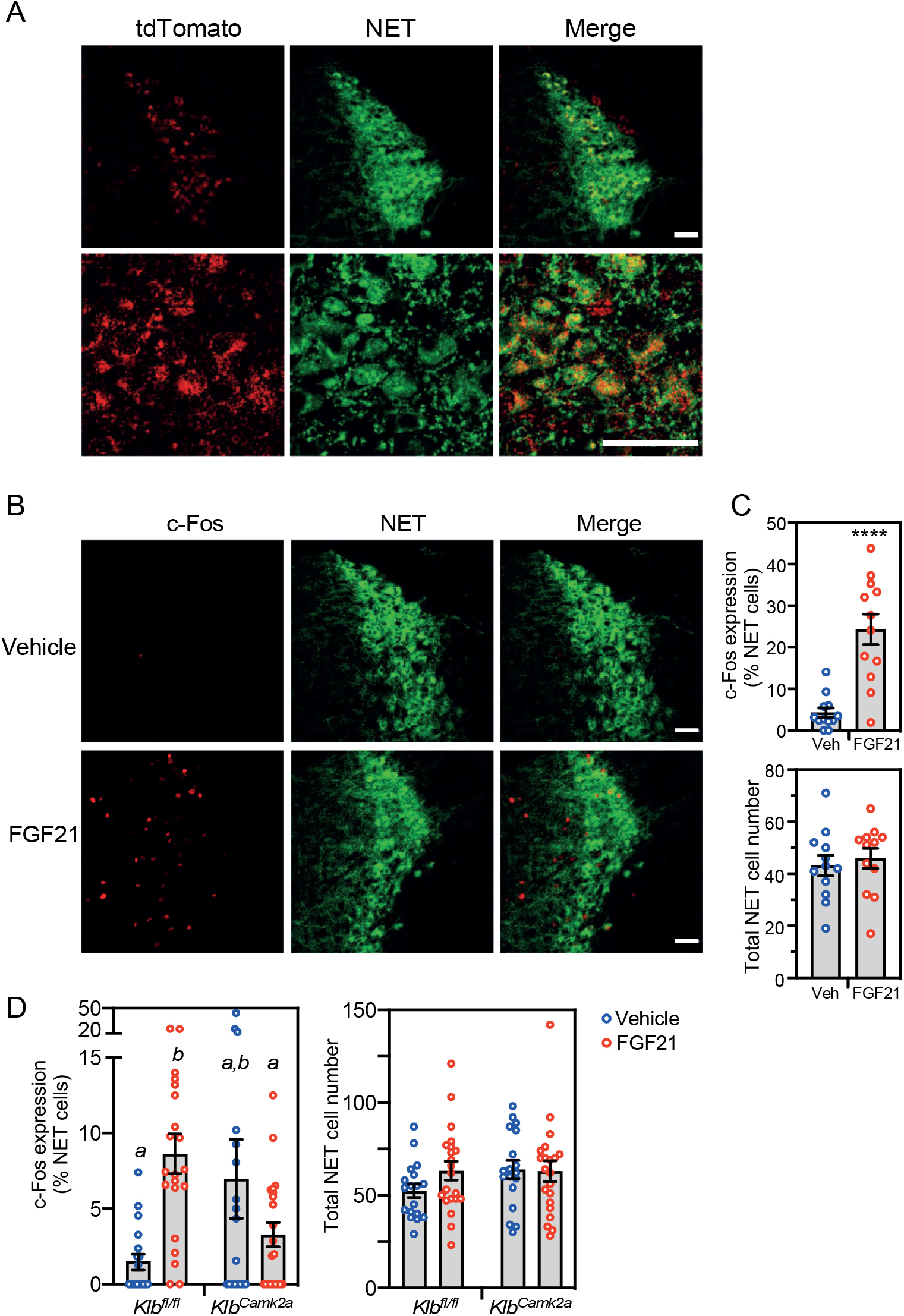
Pharmacologic FGF21 activates norepinephrine neurons in the locus coeruleus. (A) Confocal images of immunostaining performed in locus coeruleus (LC) sections prepared from mice expressing tdTomato fused to the C-terminus of KLB. Antibodies against tdTomato and norepinephrine transporter (NET) were used. Scale bars represent 50 μM. (B and C) Immunostaining for c-Fos and NET in LC sections prepared from wild-type mice treated for 2 hours with vehicle or FGF21 (1 mg/kg, i.p.). Representative confocal images are shown in (B). Scale bars represent 50 μM. Quantification of c-Fos/NET co-expression (top panel) and total NET-positive cell number (lower panel) is shown in (C) (n = 4 sections/mouse, 3 mice/group). Data represent the mean ± SEM. ****, *P* < 0.0001. (D) Immunostaining for c-Fos and NET in LC sections from groups of control (*Klb*^*fl/fl*^) and neuron-specific *Klb*^−/−^ (*Klb*^*Camk2a*^) mice treated for 2 hours with vehicle or FGF21 as in (B). Quantification of c-Fos/NET co-expression (left panel) and total NET-positive cell number (right panel) is shown (n = 3 sections/mouse, 6-7 mice/group). Data represent the mean ± SEM. Different lowercase letters indicate statistical significance (p < 0.05).

Consistent with the *Fgfr1c*/*Klb* expression data, pharmacologic FGF21 administration induced c-Fos immunoreactivity in NET^+^ LC neurons of WT mice (Figure 6B and 6C). When this same pharmacologic experiment was performed in neuron-specific *Klb*^*Camk2a*^ mice, there was no induction by FGF21 but a trend towards increased basal c-Fos expression (Figure 6D).

To test whether noradrenergic neurons are required for FGF21’s amethystic activity, we first used DSP-4, a neurotoxin that readily crosses the blood–brain barrier and selectively and irreversibly inhibits NET, thereby disrupting NE signaling (Ross and Stenfors, 2015). Mice pretreated with DSP-4 were completely refractory to FGF21’s amethystic effect on righting reflex (Figure 7A). Upon its release from neurons, NE acts on α_1-_ and β-adrenergic receptors in multiple subcortical regions to stimulate arousal (Berridge et al., 2012). Accordingly, FGF21’s amethystic effect was also blocked by the selective α_1_- and β-adrenergic receptor antagonists, prazosin and propranolol, respectively (Figures 7B and 7C). As previously reported, prazosin on its own also prolonged ethanol-induced loss of righting reflex (Figure 7B) (Malinowska et al., 1999). These pharmacologic data show that FGF21 exerts its amethystic activity by activating noradrenergic neurons in the nervous system.

**Figure 7.**
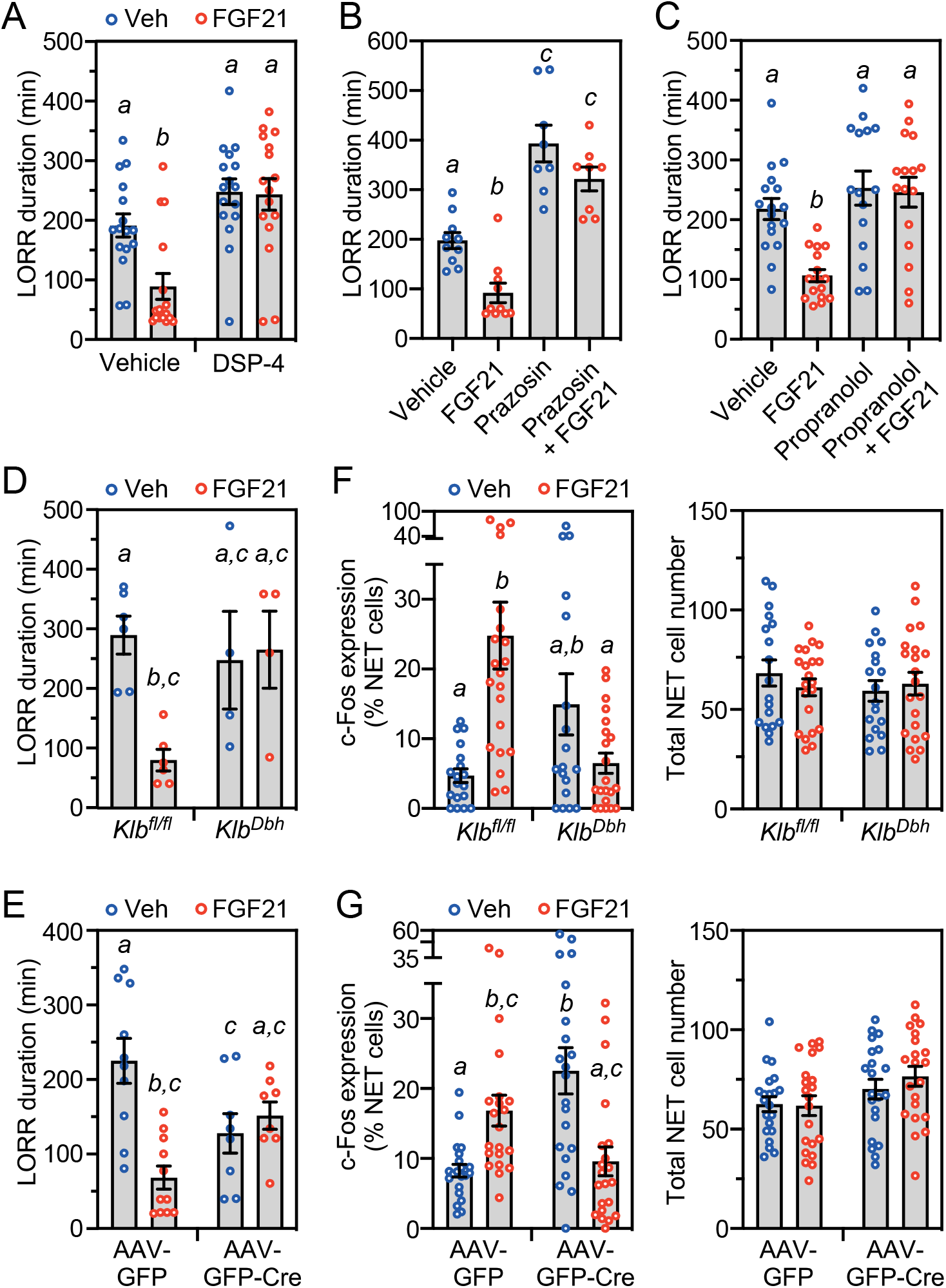
FGF21 exerts its amethystic activity through noradrenergic neurons in the locus coeruleus region. (A)Wild-type (WT) mice were injected with DSP-4 (50 mg/kg, i.p.) or vehicle. Two days later, the mice were administered ethanol (5 g/kg, oral gavage) followed 1 hour later by injection of FGF21 (1 mg/kg, i.p.) or vehicle (n = 16 mice/group). Loss of righting reflex (LORR) duration was measured after FGF21 or vehicle administration. (B)WT mice were administered ethanol (5 g/kg, oral gavage) followed 1 hour later by injection of vehicle, FGF21 (1 mg/kg, i.p.), prazosin (0.8 mg/kg) or FGF21+prazosin (n = 8-10 mice/group). LORR duration was measured after vehicle or FGF21 administration. (C)The experiment was performed as in (B) except mice were injected with vehicle, FGF21 (1 mg/kg, i.p.), propranolol (10 mg/kg, i.p.) or FGF21+propranolol (n = 15-17 mice/group). LORR duration was measured after vehicle or FGF21 administration. (D)Control (*Klb*^*fl/fl*^) and noradrenergic neuron-specific *Klb*^**−/−**^ (*Klb*^*Dbh*^) mice were administered ethanol (5 g/kg, oral gavage) followed 1 hour later by injection of vehicle or FGF21 (1 mg/kg, i.p.). LORR duration was measured after FGF21 or vehicle administration (n = 4-6 mice/group). *Klb*^*fl/fl*^ mice bilaterally injected in the locus coeruleus (LC) region with adeno-associated viruses (AAV) expressing either control (GFP) or GFP-Cre were administered ethanol (5 g/kg, oral gavage) followed 1 hour later by injection of vehicle or FGF21 (1 mg/kg, i.p.). LORR duration was measured after FGF21 or vehicle administration (n = 8-11 mice/group). (F, G) Immunostaining for c-Fos and norepinephrine transporter (NET) was performed in LC sections from groups of control (*Klb*^*fl/fl*^) and *Klb*^*Dbh*^ mice (F) or *Klb*^*fl/fl*^ mice bilaterally injected in the LC region with AAV-GFP or AAV-GFP-Cre (G) treated for 2 hours with vehicle or FGF21 (1 mg/kg, i.p.). Quantification of c-Fos/NET co-expression (left panel) and total NET-positive cell number (right panel) is shown. For (F), n = 3 sections/mouse, 6-7 mice/group. For (G), n = 7 sections/mouse, 3 mice/group. All data represent the mean ± SEM. Different lowercase letters indicate statistical significance (p < 0.05). See also Figure S2.

Lastly, we performed genetic *Klb* knockout studies to examine whether FGF21 acts directly on noradrenergic neurons in the LC region. *Klb*^*fl/fl*^ mice were crossed with *Dbh*-Cre mice to selectively disrupt FGF21 activity in noradrenergic neurons. In separate experiments, adenovirus-associated viruses (AAVs) expressing either Cre recombinase fused with green fluorescent protein (GFP) or GFP alone were bilaterally injected into the LC region. Immunostaining of brain sections with a GFP antibody showed successful targeting of the LC region (Figure S2). Consistent with the pharmacologic studies above, FGF21’s amethystic effect on righting reflex was abolished in *Klb*^*Dbh*^ mice (Figure 7D) as well as in *Klb*^*fl/fl*^ mice injected with AAV-Cre (*Klb*^AAV-*Cre*^ mice) but not control AAV-GFP (Figure 7E). Interestingly, unlike *Fgf21*^*–/–*^ and *Klb*^*Camk2a*^ mice (Figures 1C and 2E), we did not observe an increase in the time required for these knockout mice to recover their righting reflex following ethanol exposure (Figures 7D and 7E). In *Klb*^AAV-*Cre*^ mice, the recovery time was even reduced compared to control mice (Figure 7E). Accordingly, there was higher basal c-Fos expression in noradrenergic LC neurons in both *Klb*^*Dbh*^ and *Klb*^AAV-*Cre*^ mice, indicating a compensatory activation of these neurons in response to the loss of KLB. FGF21 administration did not further increase the high basal c-Fos expression in either *Klb*^*Dbh*^ or *Klb*^AAV-*Cre*^ mice (Figures 7F and 7G). Instead, there was a trend toward decreased c-Fos expression in *Klb*^*Dbh*^ mice and a significant decrease in *Klb*^AAV-*Cre*^ mice treated with FGF21 (Figures 7F and G). The reason FGF21 decreases c-Fos expression in these knockout mice is at present unknown. We conclude from these knockout studies that FGF21 exerts its amethystic effect on righting reflex by acting directly on neurons that are both noradrenergic and in the LC region.

## DISCUSSION

All fruit and nectar-eating animals, including rodents and primates, are at risk for ethanol intoxication, which increases the likelihood of injury or predation. Accordingly, they have evolved enzymatic pathways to efficiently catabolize ethanol. Interestingly, based on comparative genetic analyses of alcohol dehydrogenases, many strict herbivores and carnivores that are not exposed to alcohol appear to have lost this capability, further suggesting that ethanol is an evolutionary driver (Janiak et al., 2020). The hormone FGF21 is strongly induced in liver by both ethanol and its fermentation precursor, fructose (Desai et al., 2017; Dushay et al., 2015; Fisher et al., 2017; Søberg et al., 2018; Song et al., 2018; Zhao et al., 2015). In this report, we show that in addition to suppressing ethanol preference, stimulating water consumption and protecting against liver injury, FGF21 also protects against ethanol-induced loss of righting reflex and impairment of balance via direct effects on the noradrenergic nervous system. FGF21 does this without changing the rate at which ethanol is catabolized. Surprisingly, FGF21 does not counteract the loss of righting reflex caused by other sedatives, including ketamine, diazepam and pentobarbital. Taken together, this work reveals that FGF21 is an endogenous, ethanol-selective amethystic agent that complements the liver’s alcohol metabolizing enzymes in defending against ethanol toxicity and its potentially dangerous sequelae.

Previous studies in mice and rats showed that systemic ethanol administration acutely activates neurons in the LC (Chang et al., 1995; Kolodziejska-Akiyama et al., 2005; Thiele et al., 1997). We show that this stimulatory effect requires FGF21. Our results using liver-specific *Fgf21*^*−/−*^ mice and neuron-specific *Klb*^*Camk2a*^ mice support a model in which liver-derived FGF21 acts directly on the nervous system to counteract ethanol-induced intoxication. We also use either *Klb*^*Dbh*^ mice or the pharmacologic inhibitor DSP-4 to show that FGF21 exerts its amethystic effect by acting directly on noradrenergic neurons, which regulate arousal.

Noradrenergic neurons in the LC have been shown to be more active during periods of wake than sleep, and selective optogenetic stimulation of LC neurons elicits an immediate sleep-to-wake transition (de Lecea et al., 2012). NE stimulates arousal in part through α_1_ and β-receptors in several brain regions, including the medial septal and medial preoptic areas (Berridge et al., 2012). Accordingly, we show that FGF21’s amethystic activity is abolished genetically by selectively disrupting *Klb* expression in the LC region and pharmacologically by administering α_1_ and β-adrenergic receptor antagonists. This pathway appears to be distinct from the basolateral amygdala-initiated circuit through which FGF21 suppresses ethanol preference (Flippo et al., 2022).

Previous genetic and pharmacologic studies have established that activation of NE neurons is amethystic. Dopamine β-hydroxylase knockout mice, which are unable to synthesize NE, exhibit the same prolonged righting reflex recovery time in response to binge ethanol as we observe in *Fgf2 ^−/−^*mice. Restoration of central NE rescues this phenotype (Weinshenker et al., 2000). Likewise, drugs that suppress NE concentrations or downstream cAMP signaling also affect the degree of ethanol intoxication. For example, the tyrosine hydroxylase inhibitor, α-methyl-p-tyrosine, and the α-adrenergic receptor antagonist, phentolamine, increased ethanol-induced sleep time in mice (Blum et al., 1972; Cott et al., 1976; Masserano and Weiner, 1982). α-Methyl-p-tyrosine also increased ethanol-induced impairment of psychomotor performance and reaction time in humans (Ahlenius et al., 1973). Conversely, intracerebroventricular administration of dibutyryl cAMP antagonized ethanol-induced sedation in rats in a dosedependent manner (Cohn et al., 1975). Interestingly, impaired cAMP signaling increases the sensitivity of *Drosophila* to ethanol intoxication (Moore et al., 1998), suggesting that the mechanism underlying amethystic activity may be evolutionarily conserved.

In addition to ethanol and fructose, FGF21 is also induced by starvation and low-protein diets (Badman et al., 2007; Inagaki et al., 2007; Laeger et al., 2014). We speculate that the FGF21-NE pathway may also heighten arousal and alertness in order to increase foraging during periods of nutritional deficiency. Consistent with this, we previously showed that FGF21 increases wheel-running activity in mice during the light phase, which is highly unusual behavior for these nocturnal animals (Bookout et al., 2013). In addition to arousal, the noradrenergic nervous system impacts myriad other neuronal processes including attention, memory, perception, and motivation (Berridge et al., 2012; Poe et al., 2020). Thus, the FGF21-NE pathway may modulate a variety of cognitive and affective functions to enhance survival under stressful conditions.

In summary, FGF21 serves as an endogenous hormonal signal from liver to noradrenergic neurons in the brain to defend against ethanol-induced intoxication. Post hoc pharmacologic administration of FGF21 also markedly blunts ethanol’s effects on righting reflex and balance. These results reveal a mechanism for selectively targeting noradrenergic neurons that could prove useful for treating both the loss of consciousness and impaired mobility that occur during acute alcohol poisoning.

## ACKNOWLEDGMENTS

We thank Xiaowei Zhan for statistical analysis of the rotarod data and Genaro Hernandez for advice and experimental assistance during the early stages of this project. This work was supported by the National Institutes of Health (DK120869 to M.S., AA028473 to D.J.M. and S.A.K.), the Robert A Welch Foundation (I-1275 to D.J.M., I-1558 to S.A.K.), and the Howard Hughes Medical Institute (J.M.F. and D.J.M.). This article is subject to HHMI’s Open Access to Publications policy. HHMI lab heads have previously granted a nonexclusive CC BY 4.0 license to the public and a sublicensable license to HHMI in their research articles. Pursuant to those licenses, the author-accepted manuscript of this article can be made freely available under a CC BY 4.0 license immediately upon publication.

## AUTHOR CONTRIBUTIONS

Conceptualization, M.C., D.J.M., and S.A.K.; Methodology, M.C., M.S., W.F., A.B., L.G., and Y.Z.; Investigation, M.C., M.S., A.B., L.G., K.V., and Y.Z.; Writing, M.C., M.S., J.M.F., D.J.M., and S.A.K.; Funding Acquisition, J.M.F., D.J.M., and S.A.K.; Resources, W.F. and Y.Z.; Supervision, J.M.F., D.J.M., and S.A.K.

## METHODS

### Mouse Models

All use of mice and related procedures were approved by the University of Texas Southwestern Medical Center’s Institutional Animal Care and Use Committee. Experiments were performed with male mice unless indicated otherwise. *Fgf21*^−/−^ (Potthoff et al., 2009), *Klb*^*Camk2a*^ (Bookout et al., 2013), *Fgf21*^*Alb*^ (Song et al., 2018) and *Klb*-tdTomato reporter mice (KLB-T) (Coate et al., 2017) have been described. *Klb*^*Dbh*^ mice were generated by crossing *Dbh*-Cre mice (Jackson Laboratory, Stock No: 033951) with *Klb*^*fl/fl*^ mice (Bookout et al., 2013). For AAV injections, *Klb*^*fl/fl*^ mice were anesthetized using isoflurane anesthesia (3%-4% for induction; 1.5%-2% for maintenance) and positioned in a stereotaxic instrument (David Kopf Instruments) with a temperature controller to maintain body temperature. The skull was exposed, bregma was identified, and two small holes were drilled for bilateral injection into the LC (coordinates from lambda: ±0.9 mm mediolateral, -0.9 mm antero-posterior and -3.82 mm dorso-ventral (Paxinos and Franklin, 2019)). Dorso-ventral coordinates are relative to pia. A volume of 300 nl (1 × 10^12^ genomic particles/µl) of AAV8-GFP or AAV8-GFP-Cre virus (UNC Vector Core) was injected bilaterally into the LC. The virus was infused at 50 nl/minute using a microinjection syringe pump and Micro2T controller system with a 34G Nanofil needle (World Precision Instrument, UMP3T-1). After each injection, the needle was maintained in position for 10 minutes to prevent backflow and then slowly removed over 5 minutes. The skin was closed using sutures. Mice were allowed to recover for two weeks before use in experiments.

*Fgf21*^−/−^, *Fgf21*^*Alb*^, and KLB-T mice were on a C57BL/6J background and *Klb*^*Camk2a*^ and *Klb*^*Dbh*^ mice were on mixed C57BL/6J;129/Sv backgrounds. C57BL/6J mice were generated by the University of Texas Southwestern Medical Center animal breeding core or purchased from Jackson Laboratory. Mice were housed in a temperature-controlled environment with 12 hour light/dark cycles and fed standard rodent chow ad libitum. All experiments were performed on age- and sex-matched mice. Recombinant human FGF21 protein was generously provided by Novo Nordisk.

### Loss of Righting Reflex (LORR) Studies

Time to LORR was defined as the time between ethanol administration and LORR. Once ataxic, mice were placed in a supine position in V-shaped plastic troughs and the time measured until they were able to right themselves three times within 30 seconds, which was defined as the duration of LORR. For studying FGF21 effects on ethanol-induced LORR, mice were administered ethanol (5 g/kg) by oral gavage followed 1 hour later by i.p. injection of either FGF21 or vehicle. For the anesthetics study, ketamine (200 mg/kg), diazepam (30 mg/kg), pentobarbital (55 mg/kg) or ethanol (4.3 g/kg) were i.p. injected followed 30 minutes later by i.p. injection of FGF21 or vehicle. Diazepam and ketamine were diluted in 0.9% saline and pentobarbital was dissolved in 0.9% saline containing 10% ethanol.

### DSP-4, Prazosin and Propranolol studies

For the DSP-4 studies, C57BL/6J mice were i.p. injected with either DSP-4 (50 mg/kg) or vehicle. Two days later, mice were administered ethanol (5 g/kg) by oral gavage followed 1 hour later by i.p. injection of either FGF21 or vehicle. For the prazosin and propranolol studies, mice were administered ethanol by oral gavage followed 1 hour later by i.p. injection of FGF21 or vehicle in the presence or absence of prazosin (0.8 mg/kg) or propranolol (10 mg/kg). DSP-4, prazosin and propranolol were all dissolved in 0.9% saline just prior to use.

### Rotarod Studies

Mice were trained on a Rotamex-5 rotarod (Columbus Instruments) spinning at 5 rpm, with training complete when mice were able to stay on the rotarod for 60 seconds. To evaluate FGF21’s effect on ethanol-impaired motor coordination, mice were i.p. injected with ethanol (2 g/kg) followed 30 minutes later by i.p. injection of either FGF21 or vehicle. Time on the rotarod, with a maximum of 60 seconds, was measured at regular intervals.

### c-Fos Induction and Immunohistochemistry

For the c-Fos induction studies, mice were habituated for four days by either i.p. injection of 0.9% saline or oral gavage with water. On the fifth day, mice were i.p. injected with vehicle or FGF21 (2 hour treatment), or orally gavaged with water or ethanol (5 g/kg) as indicated in the figure legends. Mice were anesthetized with isoflurane and transcardially perfused first with PBS followed by 10% neutral buffered formalin (NBF). Brains were fixed for 24 hours in 10% NBF at 4°C and slices were prepared using a Leica VT1000S vibratome at a thickness of 50 µm. Brains from untreated KLB-T mice were processed in the same way. Slices were incubated for 1 hour in blocking buffer (1% bovine serum albumin, 5% normal goat serum, 0.3% Triton X-100 in PBS) at room temperature with shaking followed by incubation in primary antibodies, including antibodies against NET (Mab Technologies, 1:1000 dilution), cFos (Cell Signaling Technology, 1:1000 dilution), red fluorescent protein (Rockland, 1:500 dilution) and GFP (Aves Labs, 1:2000 dilution) for 48 hours at 4°C. Free-floating slices were washed 3 times in PBS for 10 minutes followed by incubation for 1 hour at room temperature with Alexa Fluor-conjugated secondary antibodies, including goat anti-mouse, goat anti-chicken, goat anti-rabbit IgGs (Invitrogen, 1:500 dilution), and DAPI (Fisher Scientific, 1:5000 dilution) in blocking buffer.

Slices were washed 3 times for 10 minutes in PBS and mounted with Aqua-Poly/Mount (Polysciences). Images were taken using a Zeiss LSM780 confocal microscope and images were processed using ImageJ software. c-Fos counts were performed blinded.

### In Situ Hybridization Analysis

C57BL/6J mice were anesthetized with isoflurane and transcardially perfused first with PBS followed by 10% NBF. Brains were fixed for 24 hours at 4°C in 10% NBF and then switched to 30% sucrose for 24 hours. Brain slices (25 µm) were cut using a freezing microtome (Leica), collected in PBS and treated with hydrogen peroxide for 10 minutes. After a PBS rinse, slices were mounted and desiccated overnight at room temperature. In situ hybridization was performed using RNAScope multiplex fluorescence kits (cat# 323110) and *Klb* (cat# 415221 Mm-Klb) and *Fgfr1c* (cat# 454941-C2 Mm-Fgfr1-O1-C2) probes purchased from Advanced Cell Diagnostics. Hybridized slides were incubated with amplification reagents and Opal 570 and 690 dyes (Akoya Biosciences, 1:1500 dilution) followed by sequential incubation with tyrosine hydroxylase (Aves Labs, 1:1000 dilution), biotinylated anti-chicken secondary (Jackson ImmunoResearch, 1:1000 dilution) and streptavidin AlexaFluor488 (Invitrogen, 1:1000 dilution) antibodies. Slides were rinsed, dehydrated, cleared, and mounted with EcoMount (BioCare Medical). Images were taken using a Zeiss LSM880 confocal microscope. ∼

### FGF21 and Ethanol Measurements

For measuring FGF21 and ethanol concentrations in murine plasma, blood was centrifuged at 3,000 rpm for 15 minutes immediately after collection and plasma was stored at -80°C until analysis. Plasma FGF21 concentrations were measured using an FGF21 mouse/rat ELISA kit (BioVendor) according to the manufacturer’s instructions. Plasma ethanol concentrations were measured using an EnzyChrom ethanol assay kit (BioAssay Systems) according to the manufacturer’s instructions. For measuring brain ethanol concentrations, brains were removed, frozen immediately in liquid nitrogen and stored at -80°C. Frozen whole brains were homogenized in 0.1N HCl and centrifuged at 13,000 rpm for 30 minutes at 4°C. Ethanol concentrations in the supernatants were measured using an EnzyChrom ethanol assay kit.

## Statistical Analyses

All data are expressed as the mean ± SEM. Statistical analyses were performed using GraphPad Prism Software Version 9.0. Unpaired two-tailed student’s t tests were used for two group analyses. Multiple groups were tested by one-way or two-way ANOVAs with Tukey’s multiple comparison test. For the rotarod analyses, a linear mixed effect model was fitted using the R package lme4 and group effect *P* values derived using a likelihood-ratio test. In all analyses, a *P* value < 0.05 was considered significant.

---

**Figure S1.**
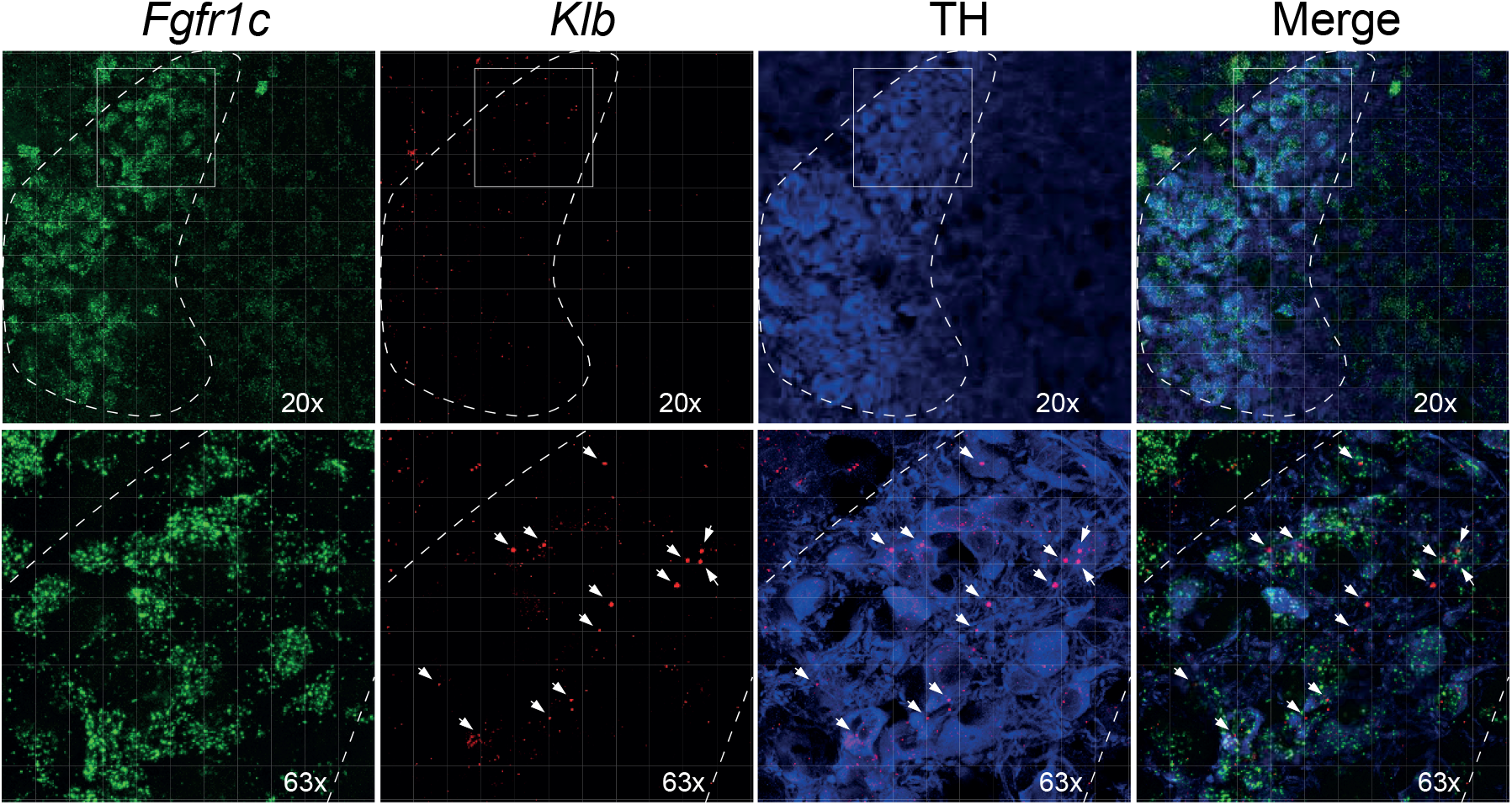
related to Figure 6. In situ hybridization of *Fgfr1c* and *Klb* mRNAs, tyrosine hydroxylase (TH) immunostaining and the merge of the three. Two different magnifications are shown, with the boxed area in the top panels expanded in the bottom panels. The fluorescent signals for *Fgfr1c* and *Klb* and are highlighted in green and red, respectively. Dotted lines outline the locus coeruleus and arrows indicate strong *Klb* signal.

**Figure S2.**
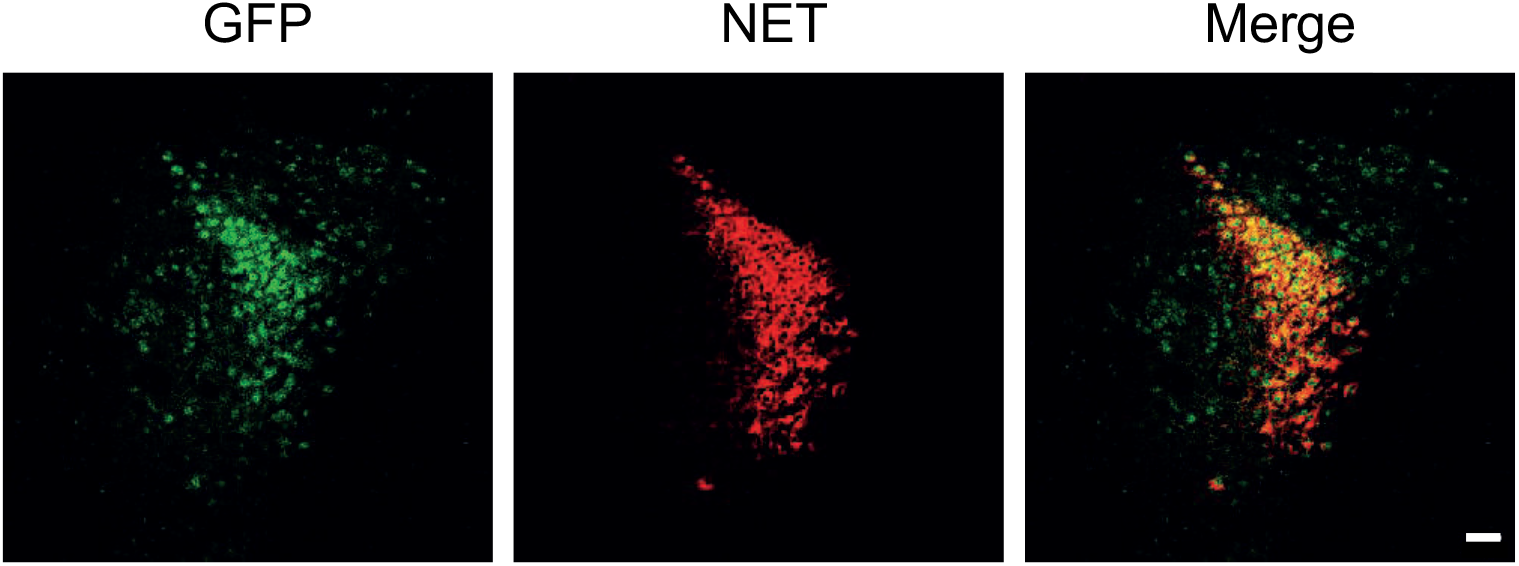
related to Figure 7. Representative confocal image of immunostaining for GFP-Cre and the norepinephrine transporter (NET) in a mouse stereotaxically injected in the locus coeruleus region with AAV-GFP-Cre. Scale bar represents 50 µM.

